# Purely enzymatic incorporation of an isotope-labeled adenine into RNA for the study of conformational dynamics by NMR

**DOI:** 10.1101/2022.02.16.480708

**Authors:** Hannes Feyrer, Cenk Onur Gurdap, Maja Marušič, Judith Schlagnitweit, Katja Petzold

## Abstract

Solution NMR spectroscopy is a well-established tool with unique advantages for structural studies of RNA molecules. However, for large RNA sequences, the NMR resonances often overlap severely. A reliable way to perform resonance assignment and allow further analysis despite spectral crowding is the use of site-specific isotope labeling in sample preparation. While solid-phase oligonucleotide synthesis has several advantages, RNA length and availability of isotope-labeled building blocks are persistent issues. Purely enzymatic methods pose as an alternative and have been presented in the literature. In this study, we report on a method in which we exploit the preference of T7 RNA polymerase for nucleotide monophosphates over triphosphates for the 5’ position, which allows 5’-labeling of RNA. Successive ligation to an unlabeled RNA strand generates a site-specifically labeled RNA. We show the successful production of such an RNA sample for NMR studies, report on experimental details and expected yields, and present the surprising finding of a previously hidden set of peaks which reveals conformational exchange in the RNA structure. This study highlights the feasibility of site-specific isotope-labeling of RNA with enzymatic methods.

## Introduction

Nuclear magnetic resonance spectroscopy (NMR) is a versatile tool to study the structure and dynamics of RNA at high resolution(1–4). Unfortunately, the chemical shift dispersion, a valuable descriptor of structure, is rather low due to high homogeneity of the helical secondary structure, and the presence of only four different building blocks. Therefore, resonances of larger RNAs often overlap severely and make resonance assignment and downstream analysis, e.g. conformational dynamics, more challenging or even impossible. Besides solutions provided by NMR itself (e.g. multidimensional experiments or selective magnetization transfer(5–7)), the issue can be tackled on the sample preparation side. Site-specific isotope labeling is one way to overcome the signal overlap that has been addressed extensively in the literature(8,9).

The most common method for site-specific incorporation of ^13^C or ^15^N-labeled nucleotides is solid-phase oligonucleotide synthesis (SPOS). Since these building blocks are often synthesized in-house, they can be modified, atom-specifically labeled or both(9,10). While SPOS has unique strengths, the most important one being simple incorporation of chemical modifications or isotope-labeled residues, the yield drops rapidly for larger constructs (>50 nucleotides (nt)). Furthermore, modified or isotope-labeled building blocks are often not commercially available or come at a significant cost.

Methods for site-specific incorporation of isotope-labeled nucleotides with *in vitro* transcription (IVT) have been presented, and can be divided in chemo-enzymatic approaches(11,12) and pause-restart methods(13). The former relies on chemical synthesis of the isotope-labeled nucleotide analog, which is then ligated to the 3’ terminus of an unlabeled strand. Another unlabeled strand is ligated to the now 3’-labeled RNA molecule(12). Liu and coworkers developed the position-selective labeling of RNA (PLOR) method, which is a sophisticated adaption of pause-restart peptide synthesis methods to RNA synthesis on agarose beads(13). Both these methods were shown to be versatile and robust, at the only caveat of needing either chemical synthesis of labeled nucleotide analog or a robotic platform for automated PLOR synthesis and modified DNA templates.

Instead of single nucleotides, shorter fragments can be labeled site-specifically and then be ligated to a larger unlabeled RNA fragment(s), which still reduces signal overlap significantly. Duss et al. presented a cut-and-paste protocol, based on *in-vitro*-transcribed labeled and unlabeled strands. These strands are then cleaved with RNase H and internal ribozymes, so that labeled fragments can be ligated to unlabeled ones, which yields a long RNA, with specific isotope-labeled segments(14,15).

Purely enzymatic methods to incorporate site-specific NMR labels and modifications have often been exploiting the fact that T7 RNA polymerase (T7RNAP) favors nucleotide monophosphates (NMP) over nucleotide triphosphates (NTP) in the first position, even though the enzyme will use NTPs if no NMPs are present. The lack of the pyrophosphate as a leaving group makes it impossible for the NMP to be incorporated into any other position than the 5’ end, making it highly specific for the 5’ position. This approach was used by Brown and coworkers to incorporate protonated GMP in the first position of a 232 kDa RNA, while all other nucleotides remained deuterated, and thus invisible in NMR spectroscopy(16). Lebars and colleagues incorporated a 6-thioguanosine into the 5’ of an RNA, which was later ligated with an unmodified RNA strand to yield a site-specifically modified RNA. This thioguanosine was later click-labeled with a proxyl-derivate radical, allowing for paramagnetic NMR studies(17).

This article uses a similar workflow, which incorporates a site-specific isotope-labeled nucleotide into an RNA molecule using only established methods based on T7 *in vitro* transcription (IVT). Our approach is similar to the method published by Lebars and coworkers and therefore strengthens the general validity of such site-specific RNA labeling methods. Importantly, compared to many other studies, we use adenosine-monophosphate (AMP) instead of guanosine-monophosphate (GMP)(16,17), which is known to hamper transcription initiation and result in lower yield(18). We thoroughly report on the sample preparation details, quantitative yields and highlight limitations and bottlenecks of the methods. Additionally, this work provides the feasibility of site-directed RNA isotope labeling to the wider NMR and RNA communities and shows how useful it can be to circumvent experimental issues with resonance overlap.

In brief, we utilize T7 RNA polymerase to incorporate a ^13^C/^15^N-labeled AMP in the 5’ position of a 16 nt RNA fragment, to then site-specifically label a longer RNA after ligation of a second RNA strand with T4 DNA ligase. We present the successful incorporation of a single ^13^C/^15^N-labeled adenosine into the 46 nt RNA molecule, which was not possible to be resonance-assigned using uniform labeling. The isotope label aids in the unambiguous assignment of the A31 residue, and further analysis of the labeled nucleotide with solution-state NMR spectroscopy gave unexpected insights into the conformational dynamics of the 46-mer.

## Material and methods

All RNA molecules were prepared using T7 IVT from a single-stranded template (double-stranded in the promoter region), which were 2’-OMe modified at the last two nucleotides of the template strand(19,20). Transcription products of similar length were purified using ion-exchange (IE) HPLC before further processing.

### RNA Sequences (figure 1)

**Figure 1:**
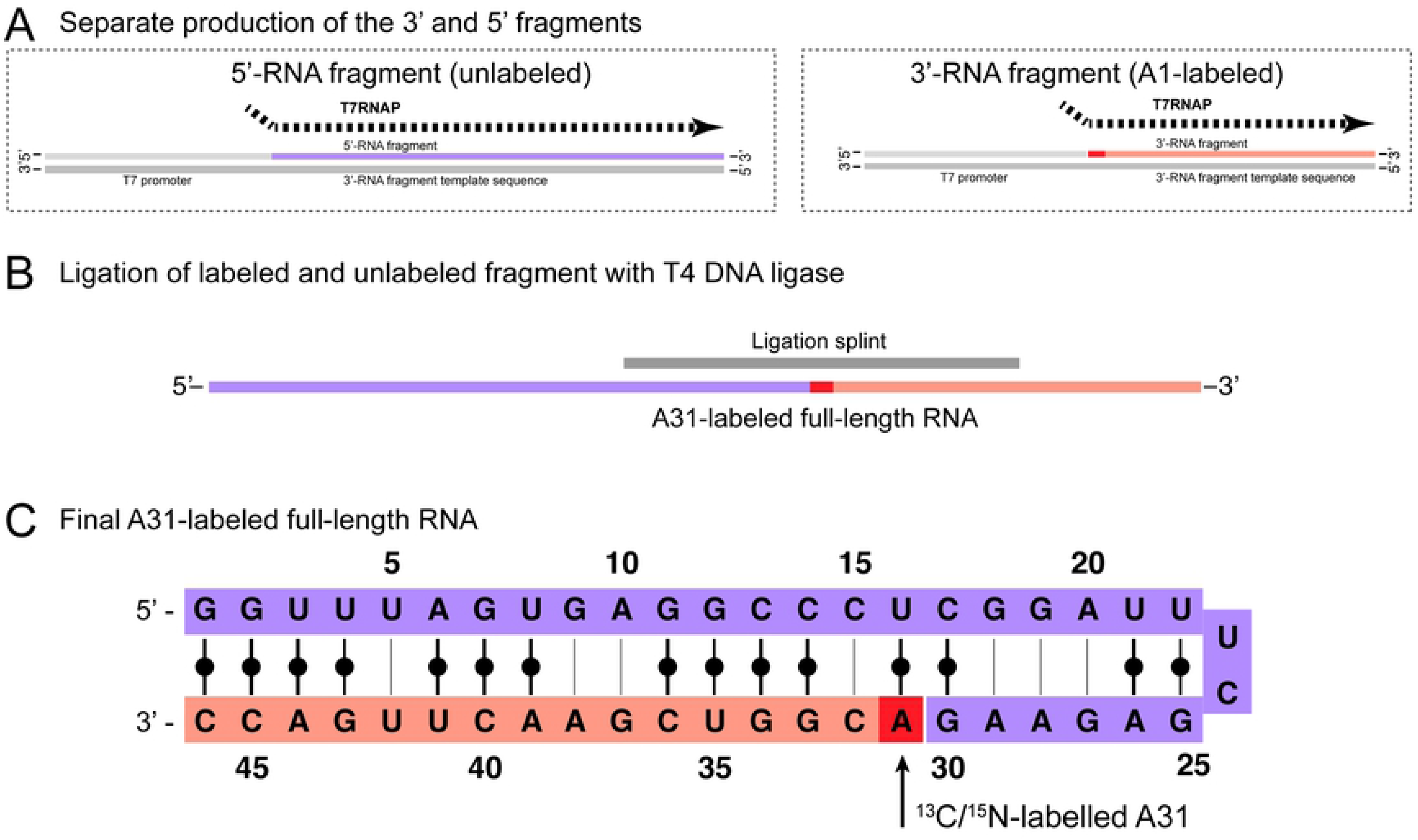
Scheme illustrating the workflow of A31-labeling of target RNA (46 nt). A: Separate production of two RNA fragments. The 3’-RNA fragment (orange) starts with the labeled nucleotide A31, which is supplied as ^13^C/^15^N-AMP into the in vitro transcription reaction. B: Ligation of unlabeled 5’-RNA (purple) and A1-labeled 3’-RNA fragment (orange) using T4 DNA ligase to produce the A31-labeled full-length RNA construct. C: Final A31-labeled RNA construct comprised of unlabeled 3’-RNA (purple) and A1-labeled (orange).

Full-length RNA (A31 in bold):

5’-GGU UUA GUG AGG CCC UCG GAU UUC GAG AAG **A**CG GUC GAA CUU GAC C −3’

5’-RNA fragment:

5’-GGU UUA GUG AGG CCC UCG GAU UUC GAG AAG −3’

3’-RNA fragment:

3’-ACG GUC GAA CUU GAC C −3’

### Template preparation

DNA templates for 5’-RNA fragment and 3’-RNA fragment (see figure 1) were ordered from IDT as custom DNA oligos with the ‘Standard Desalting’ purification option. Received oligos were diluted to 100 μM in MilliQ H_2_O and mixed by pipetting.

Template sequences with promoter complement indicated in italics are shown below:

Full-length DNA template:

5’-mGmGT CAA GTT CGA CCG TCT TCT CGA AAT CCG AGG GCC TCA CTA AAC C*TA TAG TGA GTC GTA TTA* −3’

5’-DNA fragment template:

5’-mCmUT CTC GAA ATC CGA GGG CCT CAC TAA ACC *TAT AGT GAG TCG TAT TA* −3’

3’-DNA fragment template:

5’-mGmGT CAA GTT CGA CCG T*TA TAG TGA GTC GTA TTA* −3’

T7 promoter: 5’-TAA TAC GAC TCA CTA TA −3’

T7 promoter and respective template oligo were annealed by mixing both DNA strands to a final concentration of 25 μM each and heating to 95°C for 5 minutes and successive cooling at room temperature for 30 minutes.

### T7 In vitro transcription

The MgCl_2_ (10-50 mM), Tris-Cl (60-200 mM), DTT (10 – 60 mM), Spermidine (2-20 mM), NMP (0-20 mM), NTP (3-8 mM) and DMSO (0-20%) concentrations for transcription were optimized in small scale reactions (50 μL) by rational variation and analyzed on 20% denaturing PAGE (data not shown). Conditions giving the strongest target band were then scaled up for purification and NMR sample preparation (see table 1). Reagents were added in order as shown in the table 1.

**Table 1:**
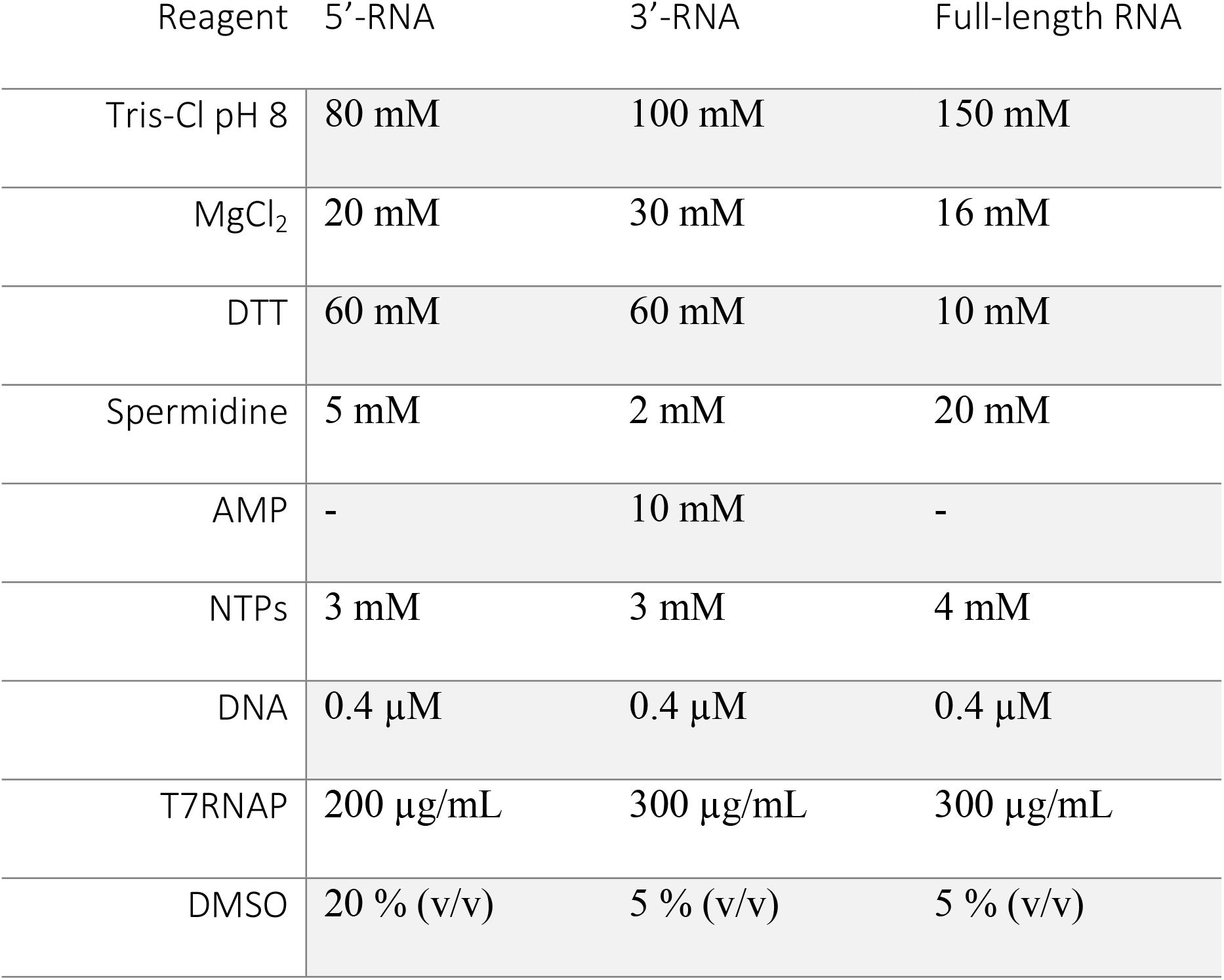
Optimized IVT conditions for RNA samples

| Reagent | 5'-RNA | 3'-RNA | Full-length RNA |
| --- | --- | --- | --- |
| Tris-Cl pH 8 | 80 mM | 100 mM | 150 mM |
| MgCl<sub>2</sub> | 20 mM | 30 mM | 16 mM |
| DTT | 60 mM | 60 mM | 10 mM |
| Spermidine | 5 mM | 2 mM | 20 mM |
| AMP | - | 10 mM | - |
| NTPs | 3 mM | 3 mM | 4 mM |
| DNA | 0.4 µM | 0.4 µM | 0.4 µM |
| T7RNAP | 200 µg/mL | 300 µg/mL | 300 µg/mL |
| DMSO | 20 % (v/v) | 5 % (v/v) | 5 % (v/v) |

T7RNAP has been expressed in *E.coli* and purified by His-tag affinity purification and size-exclusion chromatography by the protein science facility (PSF) at the Karolinska Institute.

### Denaturing polyacrylamide gel electrophoresis

All analytical gels were performed in 20% polyacrylamide gels (acrylamide:bisacrylamide 19:1) using 8×12 cm plates in a mini-PROTEAN system (BioRad). Acrylamide solution was prepared in 1X TBE buffer (89 mM Tris, 89 mM Boric acid, 2 mM EDTA, pH 8.3) and 8 M urea. 5 mL acrylamide solution were polymerized in a 15 mL centrifuge tube by adding 4 μL N,N,N,N-tetramethylethylenediamine (TEMED) and 40 μL 10% ammonium persulfate (APS) solution (v/w). The solution was mixed by inverting before pouring between the prepared glass plates (0.75 mm spacer). Gels were run in 1X TBE buffer at 350 V for typically 60 – 75 minutes. The wells were cleaned from urea by flushing with running buffer using a syringe before loading the sample.

1 μL sample was diluted in 9 μL loading solution (5 mM EDTA, 300 μM bromophenol blue in formamide) and heated to 95 °C for 2 minutes. Typically, 1 μL of the diluted sample was loaded onto the gel. Gels were stained with SYBRGold (Thermo) according to the manufacturer’s instructions and illuminated in an Amersham ImageQuant 800 UV at 498 nm excitation wavelength.

### HPLC purification

HPLC purification was performed following previously published methods(21,22). Large scale reactions (transcription or ligation) were quenched by adding EDTA to 50 mM final concentration and filtered using a 0.2 μm cellulose acetate syringe filter.

The Dionex Ultimate 3000 UHPLC system (Thermo) was equipped as follows: DNAPac PA200 22×250 Semi-Prep column, LPG-3400RS Pump, TCC-3000RS Column thermostat, AFC-3000 Fraction collector, VWD-3100 Detector. Buffer A: 20 mM sodium acetate, 20 mM sodium perchlorate, pH 6.5. Buffer B: 20 mM sodium acetate, 600 mM sodium perchlorate, pH 6.5. Buffers were degassed and filtered before use. Column temperature: 75°C. Flow rate: 8 mL/min. Injection loop: 5 mL. Pump sequences: 0 – 10 min: 0 % buffer B, 10 – 40 min: elution gradient, 40 – 50 min: 100 % buffer B, 50 – 60 min: 0 % buffer B. Elution gradients: 5’-RNA (30 nt): 22 – 30 % buffer B, 3’-RNA (16 nt): 15 – 25 % buffer B, Full-length RNA (46 nt): 15 – 45 % buffer B.

Collected fractions were tested for RNA of interest by loading 0.1 μL on a 20 % denaturing PAGE.

### RNA ligation

Ligation splints were obtained from IDT as standard DNA oligos, with the ‘Standard Desalting’ purification option. Received oligos were diluted to 100 μM in MilliQ H_2_O and mixed by pipetting.

Ligation splint 21 nt: 5’-AGT TCG ACC GTC TTC TCG AAA −3’

Ligation splint 26 nt: 5’-CAA GTT CGA CCG TCT TCT CGA AAT CC −3’

RNA ligations with T4 DNA ligase were optimized in 20 μL scale before scale-up of labeled samples. Conditions found were: 50 mM Tris-HCl pH 7.5, 10 mM MgCl_2_, 1 mM ATP, 10 mM DTT, 15 μM 5’-RNA fragment, 15 μM 3’-RNA fragment, 22.5 μM ligation splint, 2.5 % PEG 4000 (w/v), 10 % DMSO (v/v), 10 u/μL T4 DNA ligase (stock 400 u/μL, NEB). No difference in yield was found between the ligation splints. Reactions were incubated for 48 hours at room temperature. Ligation reactions have been purified by IE HPLC as described above, using a gradient of 15 - 45% buffer B.

### DNase I digest

6 mL of concentrated HPLC fractions in were digested DNase I at 1 u/μL in 1X DNase buffer (Thermo) and for 15 minutes at 37°C and purified by ion-exchanged HPLC as described above.

### NMR spectroscopy

NMR samples have been folded at 10 – 50 μM in NMR buffer (15 mM sodium phosphate, 25 mM NaCl, 0.1 mM EDTA, pH 6.5) by heating to 95°C for 5 minutes and snap-cooled in a water/ice/salt mixture for 30 minutes. The sample was concentrated to 225 μL in a centrifugal filter unit (Amicon Ultra-2, 3 kDa cutoff). 10 % D_2_O (v/v) was added to reach sample volume of 250 μL and transferred to a Shigemi tube.

NMR experiments were acquired on a 600 MHz Bruker Avance III spectrometer equipped with 5 mm HCNP QXI Cryo-probe. Spectra were processed using the Bruker Topspin 3.6 and 4.0.6 software.

^1^H,^13^C-HSQC spectra for the A1-labeled 3’-RNA fragment (300 μM) were recorded with 128 indirect points and 8 scans. ^1^H,^13^C-HSQC spectra for the A31-labeled full-length RNA (230 μM) were recorded with 256 points in the indirect dimension and 200 scans, summing up to 24 hours measurement time, and with 350 indirect points and 16 scans for the A/U-labeled sample (1.3 mM). 2D F_1_-^13^C-edited(23) NOESY/EXSY and ROESY experiments of the A31-labeled full-length RNA were recorded with 114 points in the indirect dimension and 384 scans (NOESY) or 567 scans (ROESY). NOESY mixing time was 175 ms and ROESY spinlock was 8 ms at 10 kHz.

## Results

### Construct design

We chose the 46 nt construct (fig. 1) because several resonances could not be assigned in a uniformly A/U-labeled sample (assignment not shown) due to spectral overlap and missing sequential connections, one of them A31. Two new constructs were designed, called 5’-RNA fragment (30 nt) and 3’-RNA fragment (16 nt), which will carry the 5’-adenosine label from AMP-labeled IVT and be ligated to the 3’-RNA fragment, as shown in figure 1A-C.

The incorporation of a single label at the 5’ position of the 16 nt 3’-RNA fragment was successful according to NMR experiments. ^13^C/^15^N labeled AMP (at 10 mM) was used at 3.3x excess over the other NTPs (3 mM each), as it was optimized in small scale reactions. A denaturing PAGE shows a product of 16 nt after IE HPLC purification (Figure 2A, lane 2). An aromatic ^1^H,^13^C-HSQC of the A1-labeled 3’-RNA fragment showed two signals (C2 and C8), as expected for a signal originating from a single ^13^C/^15^N-labeled adenosine nucleotide (figure 2B).

**Figure 2:**
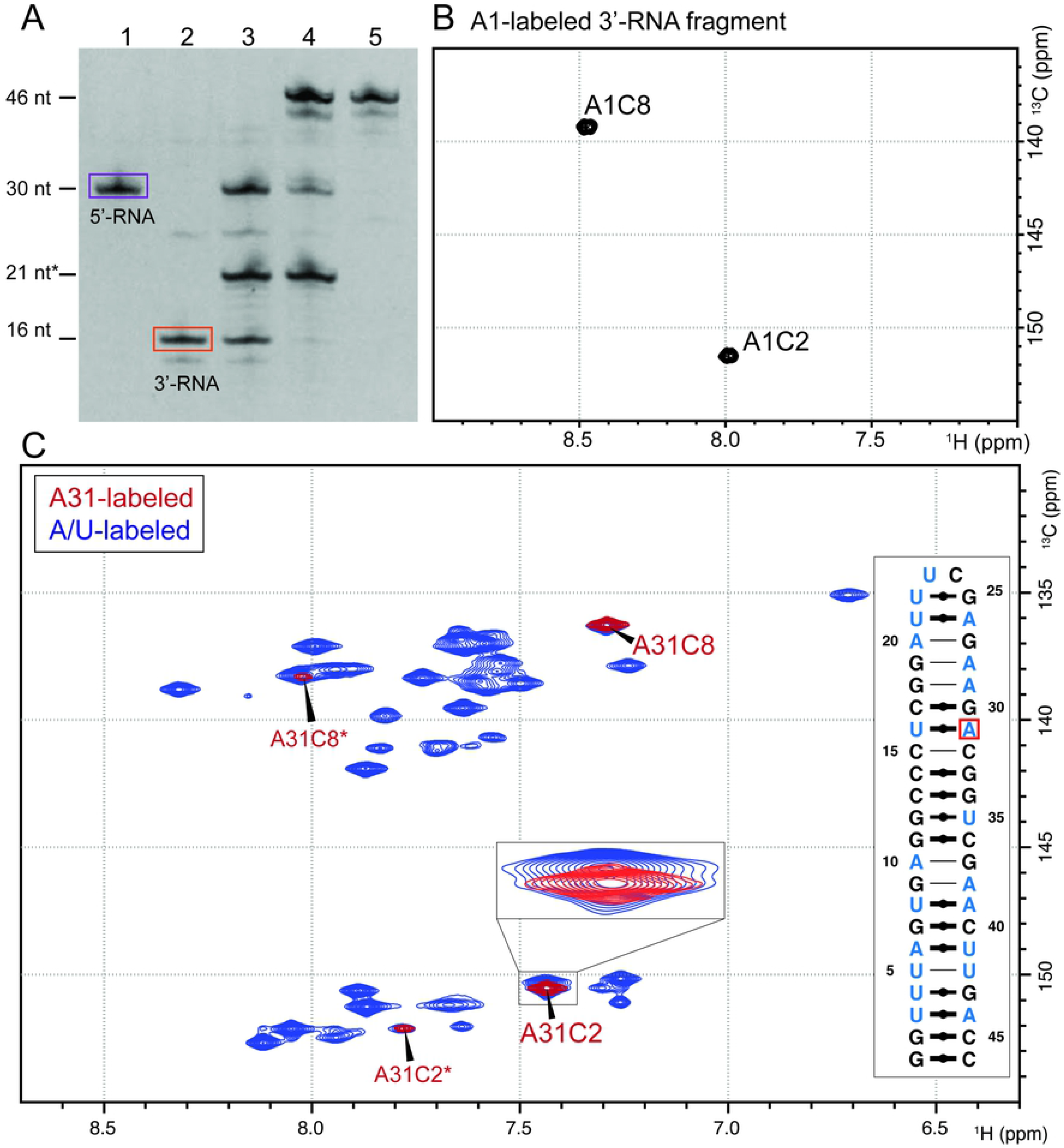
Confirmation of A31-label incorporation into the 46 nt full-length RNA. A: Denaturing PAGE of ligation reaction. Size reference nt* refers to a DNA oligo (ligation splint). Lane 1: Purified 5’-RNA fragment (purple box); Lane 2: Purified 3’-RNA fragment (orange box); Lane 3: Negative control of ligation reaction without T4 DNA ligase. Lane 4: Ligation reaction after 48 h. Lane 5: HPLC-purified full-length RNA. B: ^1^H,^13^C-HSQC of A1-labeled 3’-RNA fragment (16 nt) showing the two expected signals for C8 and C2 of A1. C: ^1^H,^13^C-HSQC spectra of the aromatic region of A31-labeled full-length RNA (red) and uniformly A/U-labeled full-length RNA (blue). For the A31-labeled RNA, two dominant set of C8/C2 signals appear alongside a weaker set of signals (A31C2* and A31C8*). The zoom shows that A31C2 does indeed overlay directly with a signal of the A/U-labeled RNA, which is partially overlapped with another signal. Furthermore, clear differences between 2B and 2C indicate proper incorporation of the isotope-labeled nucleotide into the RNA. A secondary structure prediction(25) shows which nucleotides are expected to give HSQC signals for A31-labeled RNA (red) or A/U-labeled RNA (blue).

### RNA production

For the NMR spectra shown in figures 2 and 3, an IVT reaction of the A1-labeled 3’-RNA fragment at 10 mL scale was performed. After HPLC purification, 130 nmol were obtained. This corresponds to a yield of 4.8%, whereby the limiting reagent is the most abundant nucleotide in the RNA sequence, as described previously(24).

**Figure 3:**
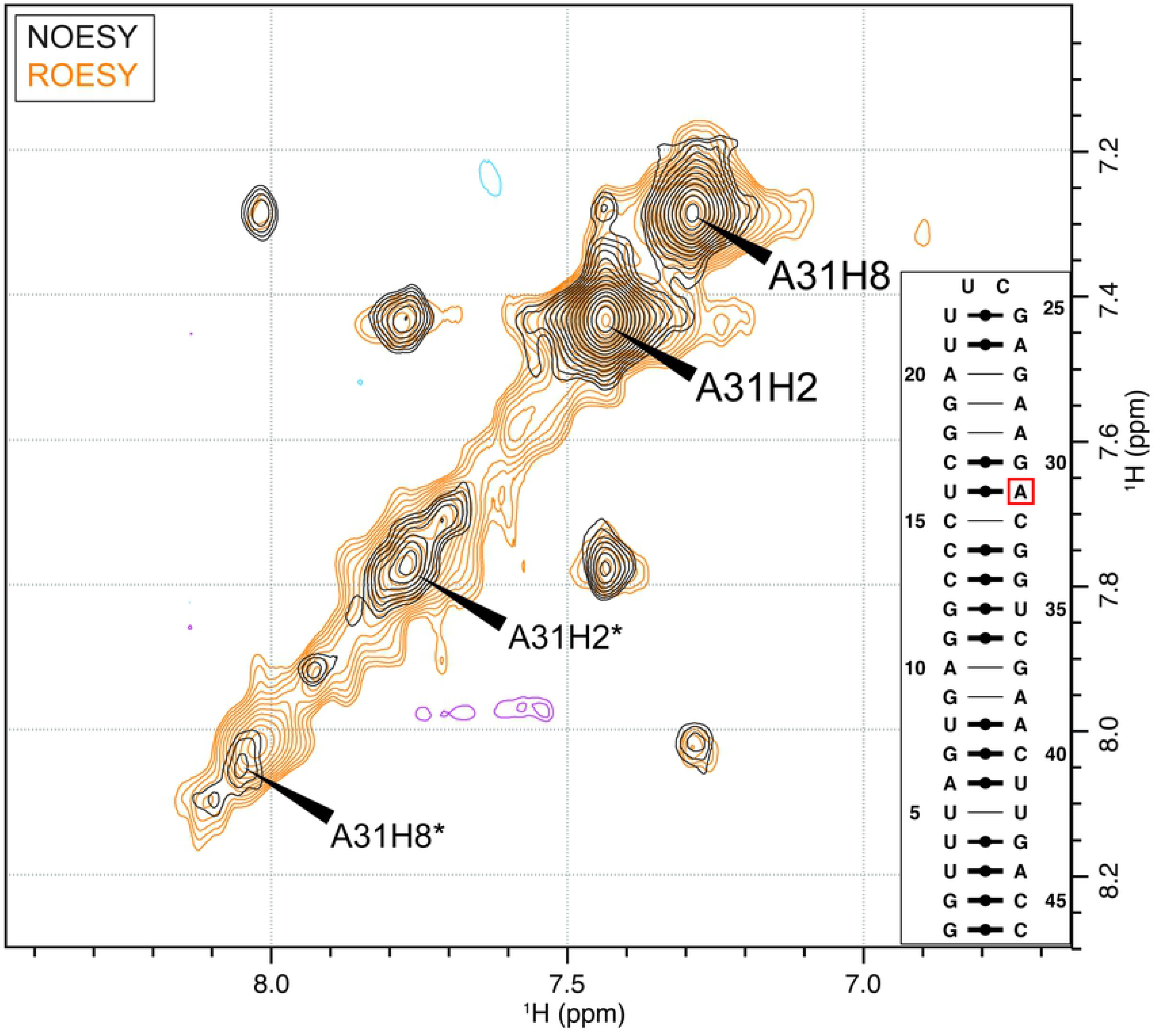
Investigation of the second set of signals in the A31-labeled RNA. ^13^C-filtered NOESY (gray/blue) and ^13^C-filtered ROESY (beige/purple) for A31-labeled RNA to probe slow conformational exchange. All cross peaks have the same sign as the diagonal signals in the respective experiment, confirming conformational exchange. A secondary structure prediction(25) is shown in the bottom right box.

The unlabeled 5’-fragment was produced at larger scale (at a yield of 5.8%) and added to stoichiometrically match the A-labeled 3’-fragment in the ligation reaction. The ligation reaction was prepared at a scale of 8.5 mL, which uses the entirety of the produced labeled 3’-RNA fragment at a final concentration of 15 μM.

The concentration of the final NMR sample was 230 μM in 250 μL, which equals 58 nmol labeled full-length RNA and implies a ligation yield, including IE HPLC purification, of 45%. In comparison, Lebars et al. and Duss et al. reported on 50 and 100% ligation yield respectively, however, using T4 RNA ligases instead of T4 DNA ligase(14,17). At this point, it should be noted that co-elution of the full-length RNA with the DNA splint required DNase I treatment and re-purification of the final sample (data not shown), which could have impacted the yield. PAGE analysis of the crude ligation reaction suggests that the entire 3’-RNA was consumed (see figure 2A, lane 4), which indicates that most of the product was lost during HPLC purification and buffer-exchange.

### Confirmation of site-specific RNA labeling

The ligation of 3’-RNA and 5’-RNA fragments was confirmed by denaturing PAGE and NMR spectroscopy. Denaturing PAGE shows a ligated band, indicating the successful formation of the 46 nt construct (figure 2A, lane 4). A second, lower band at approx. 44 nt is visible, indicating a shorter product, probably representing an impurity in the preparation of the 3’-RNA fragment (figure 2A, lane 2).

The aromatic ^1^H,^13^C-HSQC spectra of the full-length RNA sample after HPLC purification and snap-cooling, shows four peaks which overlap exactly with the A/U-labeled sample for the fully labeled construct (figure 2C). Only two signals were expected for a single adenosine in the RNA strand and figure 2B shows that only one adenosine is incorporated in the 3’-RNA fragment. However, two sets of signals each for A31C2 and A31C8 were observed. We hypothesized slow conformational exchange to be the reason for the second set of peaks., as the 5’ fragment band after HPLC purification is highly pure, as visible in Fig. 2A, lane 1.

To probe for slow conformational exchange, we performed 2D F_1_-^13^C-edited(23) NOESY/EXSY and ROESY experiments, where the signs of the cross and diagonal peaks allow the discrimination between true NOE/ROE signals and cross peaks caused by the exchange process(26). For an RNA molecule of 46 nt, NOE cross and diagonal peaks are positive in both cases. In contrast, the ROE cross and diagonal peaks are of opposite sign in the case of ‘true’ ROE peaks, but of the same sign in the case of conformational exchange. In both spectra, all cross and diagonal signals are of positive sign, indeed indicating a slow exchange process occurring (Figure 3). Conclusively, the second set of signals does not invalidate the sample preparation procedure but is a feature of the conformational dynamics of the full-length RNA, an information that would have likely not been possible to identify without the single label. At this point, the alternative conformation has not been probed further.

## Discussion

By now, methods for chemo-enzymatic(12) or purely enzymatic incorporation of site-specific spin labeling(17), including the use of NMPs for 5’-incorporation of labeled nucleotides(16) have been published. Compared to these studies, our method uses minor variations, for example, in the choice of ligase (T4 DNA ligase), or RNA purification method (ion-exchange HPLC). Furthermore, we incorporated a ^13^C/^15^N-labeled nucleotide at the specific site, while other studies used either unlabeled or modified nucleotides(12,16,17,27). Our approach is similar enough to the published literature to expect a successful incorporation of the labeled nucleotide in the right position, as we have shown here. With our work, we showcase the robustness and replicability of the method, and hopefully encourage more researchers to apply site-specific labeling where other NMR approaches prove difficult.

The optimization of several reaction steps was crucial for the success of the protocol. First, transcription conditions for all RNA constructs needed to be optimized to yield enough RNA for further reactions and NMR analysis. This was achieved by small-scale IVT reactions (50 μL) and variation of MgCl_2_, Tris-Cl, NMP, NTP, and DMSO concentrations. Ligation reactions required optimization of PEG, DMSO, temperature, and incubation time. We found no change in efficacy between a 21 nt and a 26 nt DNA splint. Due to several HPLC purification steps the final yield is rather low, yet sufficient for the assignment of the unknown nucleotide. Scale-up of the reaction seems feasible, however, one can expect diminishing returns due to the limited separation capacity of the HPLC columns with larger injection amounts. Labeled nucleotides can, in principle, be recovered and purified from the IVT buffer(28,29).

The method is expected to work at a higher yield for integration of site-specific guanosine residues, as all native T7RNAP promoters contain at least two guanosine nucleotides at the 5’ end of the transcript. Probably for this reason most previous publications used GMP at the site of labeling or modification(16,17). Despite a lower yield for a 5’-terminal adenosine, we successfully produced an NMR sample that could answer questions about the assignment and conformational dynamics. This showed that the method is feasible for starting nucleotides other than GMP, however, the yield is expected to drop further when pyrimidine bases are used(18,30,31).

Another limitation of the method is the position of the label. While T7 IVT has no upper length limit (within the size limit of solution NMR), short sequences (below 15 nt) are difficult to synthesize. Labels closer to the 3’ or 5’ termini of the full-length RNA should potentially be prepared with solid-phase oligonucleotide synthesis instead.

Alternative routes for ligation include T4 RNA ligase 1 and 2, which are known to be highly efficient, and differ with regard to their sequence and structure specificity. Also, a number of ribozymes and DNAzymes have opened new possibilities to ligate RNA in different contexts(32,33).

## Conclusion

In this report, we show another application of purely enzymatic site-specific spin labeling of RNA by IVT with isotope-labeled nucleotide monophosphate and successive ligation to an unlabeled fragment (Figure 1). Finally, we could show that the single adenine isotope label reveals a second conformation, which is in slow exchange with the ground state conformation.

The strength of the method lies in the absence of an upper size-limitation over solid-phase oligonucleotide synthesis and in a small number of robust steps needed (*in vitro* transcription and successive RNA ligation) over chemo-enzymatic methods. The weak point of the method is the sequence-dependency of incorporation efficiency of the labeled nucleotide monophosphate at the 5’-end of the labeled RNA strand. By thoroughly reporting on yields and limitations, we hope to encourage researchers to use this approach to overcome limitations from spectral crowding in NMR studies of large RNA molecules.

## Acknowledgements

We thank the Petzoldlab and especially Lorenzo Baronti for inspiring discussions and the Martin Hällberg group for the generous gift of the inorganic phosphatase. We are indebted to the protein science facility (PSF) at the Karolinska Institute for reliable production of the T7 RNA polymerase. JS acknowledges funding through a Marie Sklodowska-Curie IF (EU H2020, MSCA-IF Project No. 747446). KP acknowledges funding from the Swedish Research Council (grant number 2014-04303, and NT-2018-15), the Swedish Foundation for Strategic Research (project number ICA14-0023 and FFL15-0178), Harald och Greta Jeansson Stiftelse (JS20140009), Eva och Oscar Ahréns Stiftelse, Åke Wiberg Stiftelse (467080968 and M14-0109), Cancerfonden (CAN 2015/388, CAN 2018/715), the Karolinska Institute (KID 2016-00062), the Ragnar Söderberg Stiftelse (M91/14) and the Knut och Alice Wallenberg foundation [project grant KAW 2016.0087].

